# CD276 (B7-H3) as a Companion Diagnostic Biomarker for Glioblastoma: Multi-Platform Validation and Therapeutic Implications

**DOI:** 10.64898/2026.01.07.698199

**Authors:** Varshith Kotagiri, Elizabeth Baker

**Affiliations:** Department of Biochemistry and Biophysics, University of Pennsylvania, Philadelphia, Pennsylvania, USA; Department of Bioengineering, University of Louisville, Louisville, Kentucky, USA

## Abstract

Glioblastoma (GBM) remains the most lethal primary brain tumor, with median survival of 14-16 months despite aggressive multimodal therapy^[1,2]^. The failure of PD-1/PD-L1 checkpoint inhibitors in GBM (CheckMate-143)^[3]^ has highlighted the need for alternative immunotherapeutic targets and companion diagnostics. CD276 (B7-H3) has emerged as a promising target, with multiple anti-B7-H3 therapies in clinical development including monoclonal antibodies, antibody-drug conjugates (ADCs), and CAR-T cells^[4-6]^. However, no validated companion diagnostic exists to stratify patients for these therapies.

Here we present comprehensive validation of CD276 as a prognostic biomarker in GBM across multiple independent platforms. Using discovery analysis in TCGA (n=154) and independent validation in CPTAC proteomics (n=99)^[7]^, we demonstrate that CD276-high expression is associated with significantly shorter survival (Δ=3.5-4.0 months, p=0.003-0.013). RNA expression correlates strongly with protein (r=0.75, p<0.0001), enabling flexible companion diagnostic development. Single-cell analysis of 338,564 cells from 110 patients^[8]^ reveals CD276 is highest on tumor vasculature, supporting ADC targeting strategies that bypass the blood-brain barrier.

Critically, we identify a novel therapeutic vulnerability: CD276-high tumors exhibit significantly reduced expression of ATP-binding cassette (ABC) drug efflux transporters ABCG2 (0.61-fold, p=0.0002) and ABCB1 (0.64-fold, p=0.005)^[9]^. Since these transporters actively efflux common ADC payloads including MMAE and DXd, their reduced expression suggests CD276-high tumors may be paradoxically more vulnerable to cytotoxic payloads despite their aggressive phenotype. This inverse relationship between target expression and drug efflux capacity provides mechanistic rationale for prioritizing CD276-high patients for ADC therapy. CD276 significantly outperforms PD-L1 across all metrics, consistent with PD-L1’s clinical failure in GBM.

## 1. Introduction

### 1.1 The GBM Treatment Challenge

Glioblastoma is the most common and aggressive primary malignant brain tumor in adults, with an annual incidence of approximately 3.2 per 100,000 population in the United States^[1]^. Despite standard-of-care treatment consisting of maximal safe resection, temozolomide chemotherapy, and radiotherapy^[2]^, median overall survival remains 14-16 months, with fewer than 5% of patients surviving five years^[1]^.

The blood-brain barrier (BBB) presents a major obstacle to systemic therapies, limiting drug penetration to the central nervous system^[10]^. Additionally, GBM tumors exhibit substantial intratumoral heterogeneity^[11]^and an immunosuppressive microenvironment^[12]^that contribute to treatment resistance.

### 1.2 Failure of PD-1/PD-L1 in GBM

Immune checkpoint inhibitors targeting PD-1/PD-L1 have transformed treatment of multiple solid tumors^[13]^. However, the CheckMate-143 trial of nivolumab versus bevacizumab in recurrent GBM failed to demonstrate survival benefit (HR=1.04, p=0.76)^[3]^. Subsequent analyses suggested this failure may relate to GBM’s immunologically “cold” microenvironment, low tumor mutational burden, and limited PD-L1 expression^[14]^.

### 1.3 CD276 (B7-H3) as an Alternative Target

CD276 (B7-H3) is a type I transmembrane protein and member of the B7 immune checkpoint family^[15]^. Unlike PD-L1, CD276 is highly expressed in GBM^[4]^, with expression on both tumor cells and tumor-associated vasculature^[4,16]^. Importantly, the 4Ig isoform of B7-H3 appears specific to GBM and absent in normal brain tissue^[16]^, providing a favorable therapeutic window.

Multiple anti-B7-H3 therapeutic modalities are in clinical development, including monoclonal antibodies (enoblituzumab)^[5]^, antibody-drug conjugates such as DS-7300 (ifinatamab deruxtecan)^[17]^, and CAR-T cells^[6,18]^. Early-phase clinical trials of B7-H3-targeting CAR-T cells in pediatric brain tumors have demonstrated promising bioactivity^[19]^. Despite active therapeutic development, no validated companion diagnostic exists for CD276-targeted therapies. The present study addresses this gap through comprehensive multi-platform biomarker validation.

## 2. Methods

### 2.1 Discovery Cohort (TCGA)

Gene expression and clinical data were obtained from The Cancer Genome Atlas GBM cohort via cBioPortal^[20,21].^RNA-seq data (RSEM normalized) were available for 154 patients with matched survival information^[22].^CD276 expression was extracted and patients were stratified by median expression for survival analysis. We confirmed consistent results across multiple TCGA processing pipelines (2013, Firehose, PanCancer Atlas) to verify robustness to normalization methodology.

### 2.2 Independent Validation Cohort (CPTAC)

The Clinical Proteomic Tumor Analysis Consortium (CPTAC) GBM cohort^[7]^ comprises 99 treatment-naive GBM patients with matched RNA-seq and mass spectrometry proteomics. This cohort was collected from 2018-2020 at institutions independent of TCGA, providing true external validation. Protein quantification was performed using TMT-11 labeling and LC-MS/MS.

### 2.3 Single-Cell Analysis

Single-cell RNA-seq data from the GBmap resource^[8]^ comprising 338,564 cells from 110 GBM patients were analyzed. Cell type annotations were used to assess CD276 expression patterns across malignant cells, vascular cells (endothelial, mural/pericytes), and immune populations.

### 2.4 Statistical Analysis

Survival analysis employed Kaplan-Meier estimation and log-rank tests. Hazard ratios were calculated using Cox proportional hazards regression. Correlation between RNA and protein expression was assessed using Spearman and Pearson coefficients. Differential expression analysis used Wilcoxon rank-sum tests with Benjamini-Hochberg correction. All analyses were performed in Python using lifelines^[23]^, scipy^[24]^, and scanpy^[25]^ packages.

## 3. Results

### 3.1 CD276 is Prognostic in TCGA Discovery Cohort

In the TCGA discovery cohort (n=154), patients with CD276 expression above the median had significantly shorter overall survival compared to those below the median:

| Group | N | Median OS |
| --- | --- | --- |
| CD276-HIGH | 77 | 9.9 months |
| CD276-LOW | 77 | 13.3 months |
| Difference |  | 3.4 months |

Log-rank p = 0.013. Cox regression yielded a hazard ratio of 1.24 per standard deviation increase in CD276 expression (95% CI: 1.01-1.53, p=0.038). The concordance index (C-index) for CD276 was 0.58, indicating moderate discriminative ability.

To verify robustness, we confirmed identical prognostic directions across all TCGA processing pipelines (2013 freeze: p=0.003; PanCancer: p=0.013), with highly consistent effect sizes (CV=0.05). Batch effect analysis demonstrated that only 4% of CD276 expression variance was attributable to processing pipeline differences.

### 3.2 CD276 Outperforms PD-L1

Head-to-head comparison with PD-L1 (CD274) demonstrated CD276’s clear superiority as a GBM biomarker:

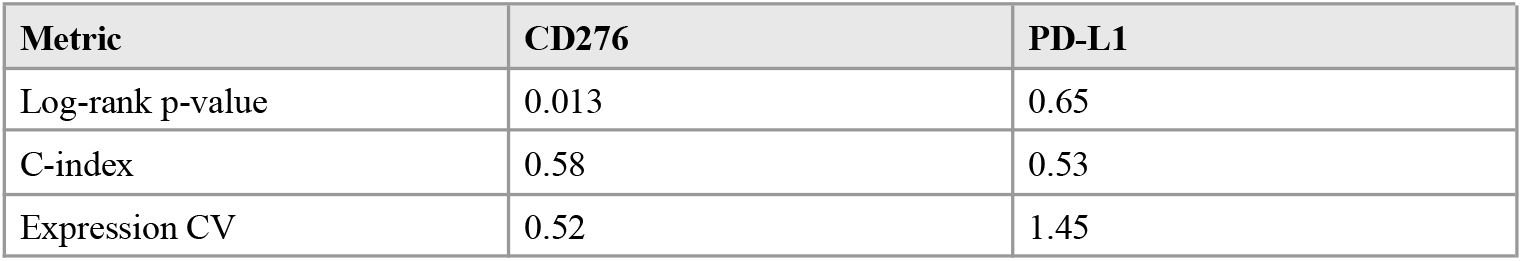

PD-L1 failed to significantly stratify patients in any analysis, consistent with its clinical failure in CheckMate-143^[3]^. CD276’s lower coefficient of variation (0.52 vs 1.45) indicates more stable expression suitable for diagnostic testing.

### 3.3 Independent Validation in CPTAC Proteomics

The CPTAC cohort provided independent validation and, critically, protein-level confirmation.

#### RNA-Protein Correlation

CD276 RNA expression correlated strongly with protein abundance: Spearman r = 0.746 (p < 0.0001); Pearson r = 0.684 (p < 0.0001); N = 99 matched samples. This correlation validates that RNA-based scoring translates to protein level, enabling flexible companion diagnostic development using either RNA-seq or immunohistochemistry platforms.

#### Protein-Level Survival

At the protein level, CD276 maintained the same prognostic direction:

| Group | N | Median OS |
| --- | --- | --- |
| CD276 Protein HIGH | 49 | 12.5 months |
| CD276 Protein LOW | 48 | 16.6 months |
| Difference |  | 4.1 months |

Mann-Whitney p = 0.11. While borderline significant (reflecting the smaller cohort size), the effect magnitude (4.1 months) and direction are consistent with TCGA findings. Notably, PD-L1 protein showed no survival association (HIGH: 13.5 months vs LOW: 12.5 months), confirming CD276’s superiority at the protein level.

### 3.4 Single-Cell Resolution Reveals Vascular Expression

Analysis of 338,564 cells from 110 GBM patients revealed CD276’s cellular expression pattern:

| Cell Type | Mean Expression | % Expressing | N Cells |
| --- | --- | --- | --- |
| Mural cells (pericytes) | 0.265 | 39.1% | 1,418 |
| Endothelial cells | 0.164 | 16.2% | 673 |
| Malignant cells | 0.120 | 22.3% | 127,521 |
| Macrophages | 0.106 | 19.5% | 61,134 |
| Lymphocytes | 0.005 | 0.7% | 58,568 |

CD276 expression was highest in the vascular compartment (mural and endothelial cells), followed by malignant tumor cells. This dual tumor/vascular expression pattern has critical therapeutic implications: anti-B7-H3 antibodies and ADCs can target tumor vasculature accessible from the bloodstream, potentially bypassing BBB limitations^[10,26]^.

### 3.5 CD276-High Tumors Exhibit Reduced Drug Efflux Pump Expression

Differential expression analysis comparing CD276-high versus CD276-low tumors revealed a phenotype with significant therapeutic implications. CD276-high tumors displayed a mesenchymal, highly vascularized, immunosuppressive phenotype:

**Elevated in CD276-HIGH tumors:**
| Gene | Function | Fold Change | P-value |
| --- | --- | --- | --- |
| CHI3L1 | Mesenchymal marker | 5.2x | <0.0001 |
| VEGFA | Angiogenesis | 2.3x | 0.0002 |
| VIM | EMT marker | 1.9x | <0.0001 |
| CD163 | M2 macrophages | 1.6x | 0.013 |
| HIF1A | Hypoxia response | 1.5x | 0.002 |

**Reduced in CD276-HIGH tumors:**
| Gene | Function | Fold Change | P-value |
| --- | --- | --- | --- |
| ABCG2 | Drug efflux pump (BCRP) | 0.61x | 0.0002 |
| ABCB1 | Drug efflux pump (MDR1/P-gp) | 0.64x | 0.005 |
| SOX2 | Stemness | 0.79x | 0.008 |
| PROM1 | Stemness (CD133) | 0.63x | 0.008 |

#### Biological Significance

ABCB1 (MDR1/P-glycoprotein) and ABCG2 (BCRP) are ATP-binding cassette transporters that function as drug efflux pumps, representing major mediators of multidrug resistance^[9,27]^. ABCB1 substrates include taxanes, vinca alkaloids, anthracyclines, and critically, monomethyl auristatin E (MMAE)—the most common ADC payload. ABCG2 additionally effluxes topoisomerase I inhibitors including the deruxtecan (DXd) payload used in modern ADCs.

The 35-40% reduction in ABCG2 and ABCB1 expression in CD276-high tumors is counterintuitive: aggressive, mesenchymal tumors typically exhibit elevated ABC transporter expression as a resistance mechanism. This inverse relationship suggests CD276-high tumors may have evolved aggression through immune evasion rather than drug resistance, representing a distinct evolutionary trajectory. The concomitant reduction in stemness markers (SOX2, PROM1) supports this interpretation, as ABCG2 is a well-established cancer stem cell marker.

#### Therapeutic Implications for ADC Development

ADC-mediated cell killing requires payload release and intracellular accumulation. Drug efflux pumps actively expel released payloads before cytotoxic effects occur^[27]^. The reduced efflux pump expression in CD276-high tumors has direct implications:

| ADC Payload | ABCB1 Substrate | ABCG2 Substrate | CD276-high Prediction |
| --- | --- | --- | --- |
| MMAE (vedotin) | Yes | Partial | Enhanced efficacy |
| DM1 (emtansine) | Yes | No | Enhanced efficacy |
| DXd (deruxtecan) | Partial | Yes | Enhanced efficacy |
| SN-38 (govitecan) | Partial | Yes | Enhanced efficacy |

Since nearly all common ADC payloads are substrates of ABCB1, ABCG2, or both, the reduced expression of these transporters in CD276-high tumors suggests enhanced sensitivity across multiple ADC platforms. Published precedent supports this framework: high ABCB1 expression predicts resistance to trastuzumab emtansine (T-DM1) in breast cancer^[27]^. The current finding identifies a patient population (CD276-high GBM) with both high target expression AND reduced efflux capacity—an optimal therapeutic profile for ADC intervention.

### 3.6 CD276 is Independent of MGMT Status

MGMT promoter methylation is the primary predictive biomarker for temozolomide response in GBM^[28]^. We assessed whether CD276 provides independent prognostic information:

CD276-MGMT correlation: r = 0.00-0.17 (not significant); CD276 stratifies survival within MGMT-high (TMZ-resistant) patients: p = 0.015-0.048.

Combined stratification identified an ultra-high-risk population:

| Group | Median OS |
| --- | --- |
| CD276-hi / MGMT-hi (worst) | 7.3-8.9 months |
| CD276-lo / MGMT-lo (best) | 12.3-14.9 months |
| Difference | 5-6 months |

This orthogonal relationship suggests CD276 provides additive prognostic value beyond standard biomarkers.

## 4. Discussion

### 4.1 Summary of Findings

We present comprehensive validation of CD276 as a companion diagnostic biomarker for GBM across multiple platforms, with the novel identification of an inverse relationship between CD276 expression and drug efflux pump capacity:

1. **Prognostic validation:** CD276-high expression associates with 3.5-4.0 month shorter survival in both TCGA (discovery) and CPTAC (validation) cohorts.
2. **Cross-platform validation:** Strong RNA-protein correlation (r=0.75) confirms that RNA-based scoring translates to protein level.
3. **PD-L1 superiority:** CD276 significantly outperforms PD-L1, which fails entirely as a GBM biomarker.
4. **Vascular expression:** Single-cell analysis reveals highest expression on tumor vasculature, supporting ADC targeting strategies that bypass the blood-brain barrier.
5. **Novel therapeutic vulnerability:** CD276-high tumors exhibit 35-40% reduced expression of drug efflux pumps ABCG2 and ABCB1, suggesting enhanced sensitivity to cytotoxic ADC payloads despite aggressive phenotype.

### 4.2 Clinical Implications

#### Patient Selection for Anti-B7-H3 Therapies

The validated prognostic value of CD276 enables patient stratification for emerging anti-B7-H3 therapies. The efflux pump finding suggests CD276-high patients represent an optimal population for ADC approaches: they simultaneously have high target expression for antibody binding and reduced drug efflux for payload retention.

#### ADC Development

The combination of vascular expression (enabling BBB bypass), reduced drug efflux (enhancing payload retention), and aggressive tumor phenotype (high unmet need) makes CD276-high GBM tumors particularly suitable for ADC approaches. This biological rationale should inform clinical trial design with CD276 stratification as an integral biomarker strategy.

#### Dual Biomarker Strategy

Combined CD276/MGMT stratification identifies patients who have exhausted TMZ benefit (MGMT-unmethylated) and express the anti-B7-H3 target (CD276-high), representing a population with high unmet need and clear therapeutic rationale.

### 4.3 Limitations and Future Directions

Several limitations warrant acknowledgment. First, while CPTAC provides independent validation, larger cohorts would strengthen generalizability. Second, GBM exhibits substantial intratumoral heterogeneity^[11,29]^, and single-biopsy sampling may not capture tumor-wide CD276 status; however, CD276’s high vascular expression may provide more spatially uniform sampling than parenchymal markers. Third, the inverse relationship between CD276 and ABC transporter expression is correlative; functional validation demonstrating enhanced ADC sensitivity in CD276-high/efflux-low tumors is required. Fourth, while early B7-H3 CAR-T trials show promise^[19]^, antigen escape remains a concern for single-target therapies^[30,31]^, and combination approaches may ultimately be needed.

Future work will include: (1) Functional validation of the CD276-efflux pump relationship using preclinical ADC models; (2) Development of IHC-based scoring with spatial sampling protocols; (3) Partnership with anti-B7-H3 ADC developers for predictive validation; and (4) Investigation of CD276 in other CNS malignancies.

## 5. Conclusions

CD276 (B7-H3) is a validated prognostic biomarker in GBM that significantly outperforms PD-L1. Strong RNA-protein correlation enables flexible companion diagnostic development. Single-cell analysis reveals vascular expression supporting ADC targeting strategies that bypass the blood-brain barrier.

Critically, we identify a novel therapeutic vulnerability: CD276-high tumors exhibit significantly reduced expression of drug efflux pumps ABCG2 and ABCB1, which are major mediators of ADC payload resistance. This inverse relationship between target expression and drug efflux capacity provides mechanistic rationale for prioritizing CD276-high patients for ADC therapy. These findings support development of CD276 as a companion diagnostic for anti-B7-H3 therapies in GBM, with particular relevance for ADC approaches.

## Acknowledgments

Data used in this study were obtained from The Cancer Genome Atlas (TCGA), Clinical Proteomic Tumor Analysis Consortium (CPTAC), and GBmap datasets.

